# ADAPTIVE MULTI-SCALE GRAPH TRANSFORMER FRAMEWORK FOR HISTOPATHOLOGICAL IMAGES

**DOI:** 10.1101/2025.10.30.685502

**Authors:** Chun-I Wu, Kalyan Banda, Elizabeth M. Swisher, Heba Sailem

## Abstract

Whole slide images (WSIs) contain hierarchical information from cellular to tissue architecture but their gigapixel scale poses major memory and computational challenges. Existing multi-scale graph and transformer models capture complex WSI features effectively but struggle with efficiency. We propose an Adaptive Multi-Scale Graph Transformer (AMGT) for WSI classification that addresses this limitation through two key modules: a Self-Guided Token Aggregation (SGTA) mechanism that fuses multi-resolution features to reduce redundancy, and a Prototypical Transformer (PT) that groups similar tokens into phenotype-representative prototypes with linear complexity. This design preserves essential spatial and semantic information, substantially lowering memory cost and improving interpretability by prototypical learning. AMGT achieves superior performance and efficiency, outperforming state-of-the-art models by 1.8% and 5.3% AUC on high-grade ovarian cancer and Camelyon16 datasets, respectively. These results demonstrate AMGT’s capacity for scalable, interpretable multi-scale representation learning.

## 1. INTRODUCTION

Whole-slide images (WSIs) are gigapixel-scale digital slides that capture intricate histological organization from cellular morphology to tissue architecture. A single 40× WSI can exceed 100,000 × 100,000 pixels, necessitating division into tens of thousands of image patches for computational modeling. Deep learning approaches, particularly weakly supervised multiple instance learning (MIL), have shown promise in predicting cancer subtypes, treatment responses, and genetic alterations [1, 2, 3, 4]. However, modeling such vast numbers of patches presents a major memory bottleneck. Transformer and graph-based methods require storing token–token or node–node correlations, where memory scales quadratically with the number of patches and number of scales incorporated. Even modest WSIs can exceed GPU limits, forcing most pipelines to either truncate spatial coverage or downsample magnifications, losing crucial cross-scale contextual cues.

### 2. RELATED WORK

Early attention-based MIL frameworks such as CLAM [5] and DSMIL [6] effectively aggregate patch-level features for slide-level prediction using lightweight attention mechanisms. While computationally efficient, these models treat patches as independent instances and lack the ability to dynamically model relationships across scales, limiting their capacity to capture hierarchical tissue organization.

To address this, transformer-based approaches such as ViT-WSI [1] introduce global self-attention to model long-range dependencies among all patches. However, the quadratic attention complexity of transformers leads to rapidly increasing GPU memory consumption as the number of patches grows, making it impractical to process full-resolution WSIs that may contain tens of thousands of tokens.

Hybrid methods such as TransMIL [7], CAMIL [8], and GTP [9] integrate convolutional or graph-based modules to improve local structure modeling. Multi-scale graph frameworks, including HIPT [10], H2-MIL [11], GRASP [12], and MS-RGCN [13], extend this idea by combining cellular- and tissue-level representations, but their dense cross-scale graph mappings incur heavy computational and memory overhead.

Recent pruning-based transformers such as LGViT [3], MSPT [2], and ZoomMIL [14] improve efficiency by clustering or discarding redundant tokens. However, without a prototypical representation to preserve recurring histological patterns, aggressive pruning often removes diagnostically important regions, leading to degraded multi-scale reasoning and reduced accuracy.

### 3. CONTRIBUTIONS

We propose the Adaptive Multi-Scale Graph Transformer (AMGT), a unified framework for accurate and memory-efficient WSI analysis. It integrates a Self-Guided Token Aggregation (SGTA) module that adaptively fuses tissue- and cell-level features by retaining salient regions, and a Prototypical Transformer (PT) that clusters similar representations into phenotype-aware prototypes with linear complexity. Together, these components enable AMGT to preserve high representational fidelity at a fraction of the memory cost. Validated on ovarian cancer and Camelyon16 datasets, AMGT sur-passes state-of-the-art methods in both accuracy and efficiency, providing a scalable and interpretable solution for large-scale histopathological image analysis.

## 4. METHODS

**Overview**. The proposed Adaptive Multi-Scale Graph Transformer (AMGT) aims to efficiently integrate finegrained cellular details with global tissue organization in WSIs under constrained GPU memory. As shown in Fig. 1a, AMGT constructs a multi-scale heterogeneous graph, fuses salient cross-scale features through a Self-Guided Token Aggregation (SGTA) module, and summarizes them using a Prototypical Transformer (PT) for slide-level prediction. This design reduces redundant computation while retaining diagnostically critical information.

**Fig. 1.**
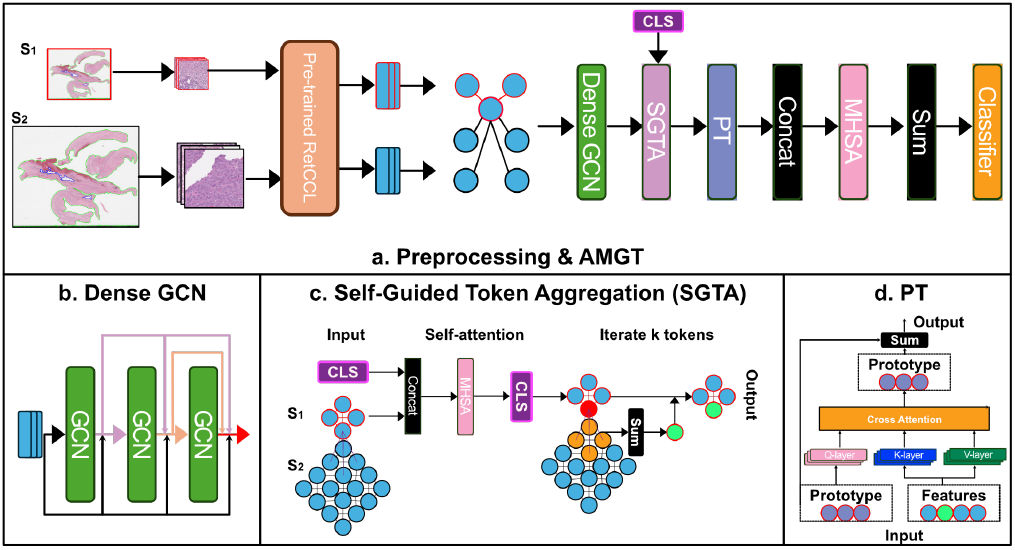
Overview of the proposed Adaptive Multi-Scale Graph Transformer (AMGT). (a) A whole-slide image is divided into low-resolution (*S*1) and high-resolution (*S*2) patches to construct a multi-scale graph. (b) Dense GCN layers propagate spatial and cross-scale features. (c) The SGTA module selects diagnostically relevant regions and fuses fine-grained features. (d) The Prototypical Transformer produces interpretable slide-level representations for classification.

### 4.1. Multi-Scale Graph Construction

Each WSI is divided into non-overlapping patches at two magnifications: S1 (5×) and S2 (10×). Background patches are removed using HSV thresholding, and remaining patches are embedded via a pretrained RetCCL encoder [3] to generate 2048-dimensional feature vectors. A heterogeneous graph *G* = (**V, E**) is then constructed, where 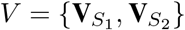 denotes node sets from each scale, and 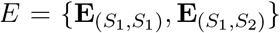defines intraand cross-scale edges based on patch centroid distances.

A Dense Graph Convolutional Network (DGCN) propagates information through this graph, allowing both local (intra-scale) and hierarchical (cross-scale) interactions while mitigating over-smoothing (Fig. 1b).

### 4.2. Self-Guided Token Aggregation (SGTA)

SGTA identifies diagnostically relevant low-resolution regions and aggregates corresponding high-resolution details (Fig. 1c). Specifically, a multi-head selfattention (MHSA) layer processes low-resolution tokens 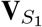 concatenated with a class token (CLS) to estimate attention weights, from which the k tokens with the highest transformer attention are selected. Each selected token sums its high-resolution neighbors from 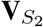 through graph-defined edges. This adaptive mechanism fuses contextual and fine-grained information, reducing token count and memory consumption while emphasizing histologically meaningful regions.

### 4.3. Prototypical Transformer (PT)

To further improve scalability, AMGT introduces a Prototypical Transformer that compresses redundant features into a compact set of learnable prototype tokens (Fig. 1d). The PT performs linear cross-attention, where prototypes query the aggregated feature sequence, capturing phenotype-level patterns with linear computational complexity. A subsequent self-attention among prototypes models inter-phenotype relations, producing a compact slide-level representation that preserves semantic diversity while minimizing memory load.

### 5 EXPERIMENTS

#### 5.1. Datasets

To evaluate the effectiveness of AMGT, we conducted experiments on in-house ovarian cancer (IHOV) for platinum response prediction and public dataset Camelyon16 [15] for breast cancer tumor classification. The IHOV dataset consists of 650 hematoxylin and eosin (H&E) stained diagnostic slides from 134 high grade ovarian cancer patients (Platinum sensitive: n=101; refractory and resistive: n=33). Camelyon16 includes 270 training WSIs (normal n=159; tumor n=111) and 129 testing WSIs (normal n=80; tumor n=49).

#### 5.2. Preprocessing

Each WSI was cut into patches of size (1024;15*×* ) and (2048; 7.5*×* ), background patches are removed by setting a threshold in HSV space and resized via BiLinear interpolation (384, 384). Each patch is then fed to the pretrained RetCCL [3] model. Graph data is constructed by considering Euclidean distance under 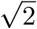 patch size from patch centroids to ensure spatial correlation between patches is preserved. Edge indices and types are used for heterogeneous graph construction.

#### 5.3. Training Strategy and Implementation

Models were trained with a fixed global seed (2025) to ensure consistent WSI feeding order across runs. The in-house ovarian cancer (IHOV) dataset was divided into 75% training and 25% testing, while Camelyon16 followed its predefined 67.5:32.5 split. All models were trained on NVIDIA A100 GPUs using half-precision (FP16). The Adam optimizer (lr = 2e-4, weight decay = 1e-5, dropout = 0.1) was used, with training capped at 200 epochs, early stopping (patience = 10), and gradient accumulation = 2 to simulate a larger batch size. The training objective combined binary cross-entropy loss with the prototype disentanglement loss proposed in [6], which helps prevent prototype collapse by encouraging each prototype to capture distinct patterns.

#### 5.4. Efficiency Evaluation

To assess computational efficiency, we use two key metrics derived from PyTorch profiling: (a) Trainable Parameters (PS), the total number of learnable weights requiring gradient updates, reflecting model complexity and storage cost; and (b) Maximum Memory Allocation (MaxMA), the peak GPU memory usage during a single training epoch on Camelyon16, accounting for both parameters and intermediate activations. MaxMA offers a realistic measure of runtime scalability, particularly critical for WSI processing where memory limits constrain batch size and resolution.

## 6. RESULTS

### 6.1 Benchmark Results

AMGT outperformed all state-of-the-art baselines with second-best memory efficiency (Table 1.). It achieved the highest AUC scores on both IHOV (0.791) and Camelyon16 (0.968), while maintaining a compact architecture. AMGT has the second smallest PS (2.34M) and MaxMA (0.277 GB), marginally larger than MSRGCN (2.01/0.196). Compared to the next-best models, ZoomMIL and TransMIL, AMGT achieved AUC gains of 1.8% and 5.3% on IHOV and Camelyon16, respectively. These findings demonstrate that AMGT can effectively integrating both local and global contextual information in a memory-efficient manner. Specifically, AMGT outperforms ZoomMIL which has a similar token pruning strategy, on both datasets, demonstrating the added benefit of explicitly modeling cross-scale relationships and phenotyping. These results highlight that accurate and compact WSI representations can be constructed without exhaustively visiting all patches across magnification levels.

**Table 1.** Benchmark comparison of AMGT with state-of-the-art models on IHOV and Camelyon16 (C16).

| Model | Params (M) | Mem (GB) | AUC-IHOV | ACC-IHOV | AUC-C16 | ACC-C16 |
| --- | --- | --- | --- | --- | --- | --- |
| ViT-WSI [1] | 18.08 | 18.08 | 0.664 | 0.771 | 0.890 | 0.853 |
| TransMIL [7] | 18.20 | 5.17 | 0.656 | 0.714 | 0.915 | 0.899 |
| DSMIL [6] | 12.59 | 0.29 | 0.740 | 0.800 | 0.891 | 0.899 |
| ZoomMIL [14] | 4.50 | 0.98 | 0.773 | 0.727 | 0.820 | 0.829 |
| MS-RGCN [13] | **2.01** | **0.20** | 0.708 | 0.771 | 0.893 | **0.907** |
| **AMGT (Ours)** | 2.34 | 0.28 | **0.791** | **0.818** | **0.968** | 0.899 |

### 6.2 Interpretability Analysis

We visualized prototype attention heatmaps to examine AMGT’s interpretability. As shown in Fig 2, while majority focused on tumor section while other prototypes attended to distinct histological regions such as adipose (p2), viable tumor nuclei–dense patches (p5), necrotic-like tissue and stromal compartments (p7), demonstrating stable and interpretable attention behavior that underscores AMGT’s ability to capture meaningful tissue structures and enhance clinical transparency.

**Fig. 2.**
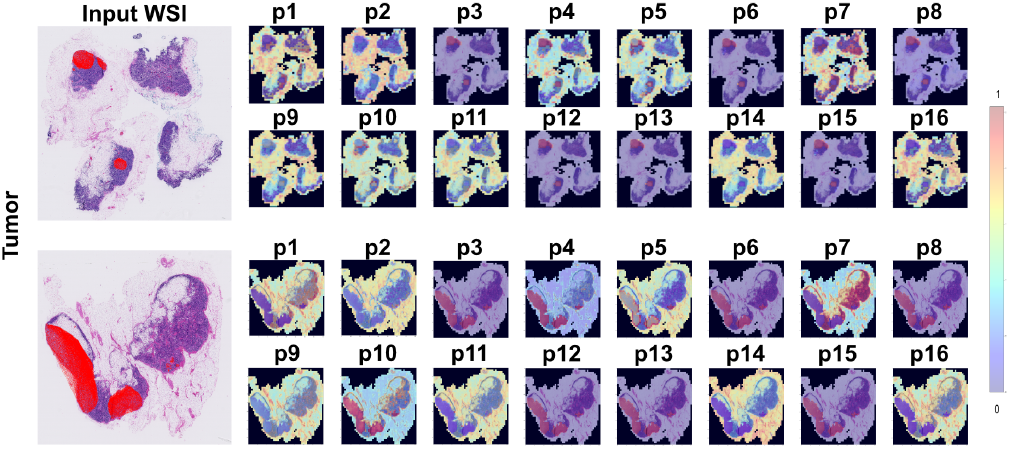
Interpretability analysis of AMGT on WSIs from the Camelyon16 dataset (test image 001 and 016). Each panel *p*_*n*_ represents the attention map of prototype n, using a jet colormap (red indicates annotated tumor regions).

### 6.3 Ablation Studies

We conducted ablation studies to evaluate the contributions of each AMGT component and its hyperparameters (Tables 2–3). Starting from a baseline graph transformer with DGCN and MHSA, we incrementally added the SGTA and PT modules. Replacing MHSA with PT improved AUC on IHOV, while SGTA alone offered limited benefits, indicating that the two modules function synergistically. SGTA identifies salient spatial regions, whereas PT compresses these features into phenotype-level prototypes; together, they achieve both higher accuracy and efficiency in multi-scale WSI classification. We examined the effects of varying the number of selected top-*k* tokens and prototype count (*p*). Moderate pruning (*k*=200) yielded the best performance, confirming that removing redundant tokens enhances learning. Increasing prototype numbers improved representation diversity, suggesting that a richer set of prototypes better captures heterogeneous tissue phenotypes. Both *k* and *p* are thus critical hyperparameters that balance efficiency and expressiveness across datasets.

**Table 2.** Ablation of AMGT components on IHOV and Camelyon16 (C16).

| Configuration | Scale | AUC-IHOV | AUC-C16 |
| --- | --- | --- | --- |
| DGCN+MHSA | 7.5×,15× | 0.67 | 0.88 |
| DGCN+PT | 15× | 0.77 | 0.75 |
| DGCN+SGTA+MHSA | 7.5×,15× | 0.67 | 0.89 |
| **AMGT (Full)** | 7.5×,15× | **0.79** | **0.97** |

**Table 3.** Effect of top-*k* tokens and prototype count (*p*) on AMGT performance.

| Setting | AUC-IHOV | AUC-C16 | $k$ | $p$ |
| --- | --- | --- | --- | --- |
| $p=16, k=100$ | 0.74 | 0.82 | 100 | 16 |
| $p=16, k=200$ | **0.79** | **0.97** | 200 | 16 |
| $p=16, k=400$ | 0.65 | 0.90 | 400 | 16 |
| $p=8, k=200$ | 0.71 | 0.76 | 200 | 8 |
| $p=4, k=200$ | 0.70 | 0.76 | 200 | 4 |
| $p=1, k=200$ | 0.72 | 0.75 | 200 | 1 |

## 7. CONCLUSION

We presented AMGT, a prototype-based multi-scale graph transformer for efficient WSI classification. By combining selective token aggregation and prototype-based compression, AMGT captures hierarchical cross-scale features with high accuracy using fewer parameters. Validated on diverse histopathology datasets, showing strong generalization and interpretability. Future work will focus on adaptive cross-scale prototypes fusion.

## ACKNOWLEDGEMENT

This work is funded by a Wellcome Career Development Award 225974/Z/22/Z.

